# Development of a Surface Programmable Activation Receptor system (SPAR): A living cell biosensor for rapid pathogen detection

**DOI:** 10.1101/687426

**Authors:** Joseph D. Kittle, Joel S. Lwande, M. Russell Williams, Richard Brody, Melissa Frenchmeyer, Jiangzhou Hua, Shengwen Liang, Kyle McQuaid, Min Mo, Allison Neese, Yuanyuan Tang, Srikanth Vedamoorthy, Lingchun Zeng, Thomas Zupancic, Charles McBrairty

**Affiliations:** Fundamental Solutions Corporation/CytoSPAR, Easton, PA; InfinixBio, Athens, OH; DevLab LLC, Cleveland, OH; David H. Murdock Research Institute, Kannapolis, NC.

## Abstract

Efficient pathogen detection is essential for the successful treatment and prevention of infectious disease; however, current methods are often too time intensive to be clinically relevant in cases requiring immediate intervention. We have developed a Surface Programmable Activation Receptor (SPAR) diagnostic platform comprised of universal biosensor cells engineered for use in combination with custom or commercial antibodies to achieve rapid and sensitive pathogen detection. SPAR cells are stably transfected Jurkat T cells designed to constitutively express a modified T cell mouse FcγRI receptor on the cell surface and a high level of the luminescent reporter protein aequorin in the cytoplasm. The modified mFcγRI-CD3ζ receptor protein binds with high affinity to the Fc region of any full-length mouse IgG2a and some IgG2 antibodies: this allows customized target detection via the selection of specific antibodies. T-cell receptor aggregation in response to target antigen binding results in signal transduction which, when amplified via the endogenous T cell signal cascade, triggers the rapid intracellular release of calcium. Increased Ca^2+^ concentrations activate the expressed reporter protein aequorin resulting in the immediate emission of detectable light. Testing demonstrates the accurate and specific detection of numerous targets including *P. aeruginosa*, *E. coli* O111, and *E. coli* O157. We report that the SPAR biosensor cell platform is a reliable pathogen detection method that enables the rapid identification of bacterial causative agents using standard laboratory instrumentation. The technology lends itself to the development of efficient point-of-care testing and may aid in the implementation of effective and pathogen-specific clinical therapies.

## Introduction

The rapid and accurate identification of causative agents is critical to the prompt application of directed, pathogen-specific antibiotic therapies. Effective and timely clinical intervention is essential for the control of infectious disease as well as in the successful treatment of bacterial infections. The observed increases in the frequency and severity of nosocomial infections (1) and the increasing prevalence of antibiotic resistance induced by non-specific antibiotic use (2) further highlight the need for informed antibiotic selection based upon precise and efficient pathogen detection.

Current bacterial identification methods include both classic procedures and novel molecular techniques. Traditional culture-based methods, while sensitive and reliable, are also labor-intensive and time-consuming and therefore often cannot provide definitive diagnostics within a clinically relevant timeframe (3). Molecular diagnostic methods include immunological assays such as ELISA (4), microarray immunoblot (5), or serological assays (6); nucleic acid-based techniques including PCR (7), DNA sequencing (8), hybridization techniques (9), or DNA/RNA microarrays (10); and mass spectrometry (11). These methods may provide increased specificity or diagnostic speed, yet often require sophisticated instrumentation, skilled personnel, and/or time-consuming sample enrichment steps, and occasionally produce false-positive or false-negative results (5, 12). Therefore, these methods are not optimal for use as rapid clinical or field diagnostics.

The SPAR cell design described in this paper takes advantage of two well-documented cellular responses and combines them in a single cell line to create a fast-acting and robust reporter platform that circumvents these difficulties.

Human T cells express peptide-specific receptors (TCRs) that engage with their cognate antigens and play a central role in the adaptive immune response (13). This extracellular recognition event triggers the activation of an intracellular network of signaling pathways that direct the transcriptional and physiologic processes responsible for T cell activation and function (14). One such aspect of signaling pathway activation is the rapid increase in intracellular Ca^2+^ concentration, attributed in part to the efflux of stored Ca^2+^ from the endoplasmic reticulum (15, 16).

Aequorin, a calcium-sensitive photoprotein isolated from the jellyfish *Aequorea victoria*, emits blue light in response to the presence of Ca^2+^ (17). The native aequorin molecule, which possesses three Ca^2+^ binding sites, is bound to its chromophoric ligand coelenterazine. Ca^2+^ binding triggers the oxidation of coelenterazine to coelenteramide, with the concomitant release of CO_2_ and the emission of detectable light (18). Recombinant expression of this protein has been exploited as a mechanism for the detection of transient changes in intracellular levels of free Ca^2+^ for decades and is well documented (19-21). Aequorin has been used as a detector of B cell activation in systems where activation of a chimeric B cell receptor leads to calcium release in response to the binding of a specific antigen (22). Cells expressing recombinant aequorin must be charged prior to use by incubation with the hydrophobic ligand coelenterazine, which readily permeates cell membranes (21).

Here we report the development of an engineered T cell line that stably expresses both a modified TCR complex and a high level of aequorin reporter protein. We demonstrate that the modified TCR complex binds with high affinity to the Fc region of the murine IgG2a isotype and to a variety of additional murine IgG2s, and is therefore capable of specific, customizable antigen binding and signal transduction. These signals, when amplified via the endogenous T cell signal cascade, result in Ca^2+^ release: local increases in cytosolic Ca^2+^ activate the SPAR cell aequorin reporter system resulting in the emission of detectable light. Thus, our results establish that the SPAR cell is a novel, self-contained biosensor platform that is effective for the rapid and sensitive identification of target antigens.

## Materials and Methods

### Experimental Design

The objective of the work described in this paper was to generate a living cell-based biosensor platform that can be readily adapted to detect any specific pathogen and deliver rapid, sensitive, and easily interpreted results. Here, we describe the methods used to generate SPAR biosensor cells and the verification of their functional capacity. We also provide data to demonstrate that SPAR cell pathogen detection is accurate and is readily customized via the addition of specific antibodies.

### Antibodies, bacterial strains, and cell lines

Goat Anti-*E. coli* O157 was purchased from KPL (Gaithersburg, MD). Mouse anti-Human IgG was purchased from Biolegend (San Diego, CA). Bovine Serum Albumin and Goat Anti-Mouse IgG HRP conjugate were purchased from Millipore Sigma (St. Louis, MO). The mouse monoclonal antibody against *P. aeruginosa* was obtained from Novus Biologicals (Centennial, CO). The following bacterial strains were purchased from ATCC (Manassas, VA): *E. coli* O111; *E. coli* O157:H7; *P. aeruginosa*; *S. enteritidis*. The mouse monoclonal antibody against *E. coli* O111 LPS was generated by Precision Antibody, Columbia, MD, as described below. The Jurkat cell line (Clone E61, ATCC® TIB 152™) was obtained from ATCC and maintained in Complete RPMI medium: RPMI 1640 (VWR, Radnor, PA) + 25 mM dextrose, 10 mM HEPES, 1 mM sodium pyruvate, 10% FBS, and 1% Pen-Strep.

### Construction of Aequorin Expression Vector pFSC005

The aequorin DNA sequence (23) was purchased (DNA2.0/ATUM, Newark, CA), amplified via PCR, restriction enzyme-digested, and cloned into the standard mammalian expression vector pEF1/myc-His B (Thermo Fisher Scientific, Waltham, MA).

### Construction of mFcγRI-CD3ζ Expression Vector pFSC048

Briefly, the CD3*ζ*SS-FcγRI-CD3*ζ* expression vector was constructed into pVitro1-Aeq vector, built in-house based on InvivoGen pVitro1-Blasti-MCS vector (InvivoGen, San Diego, CA) (unpublished data). To construct FcγRI-CD3*ζ*, the human CD3 zeta chain sequence (MyBioSource Inc, San Diego, CA) was PCR-amplified and fused to the extracellular domain of mouse FcγRI (GeneCopoeia, Rockville, MD) via overlap PCR amplification. The human CD3*ζ* signal sequence (CD3*ζ* SS) was PCR-amplified from the CD3*ζ* cDNA clone and fused to the FcγRI-CD3*ζ* PCR fragment via overlap PCR. The generated CD3*ζ* SS-FcγRI-CD3*ζ* PCR fragment was then digested and cloned into pVitro1-Aeq vector to create plasmid pFSC048.

### Generation of SPAR Jurkat P5G7 Cells

#### DNA Linearization and Purification

Plasmids pFSC005 containing the hEF-1α-Aeq construct for aequorin expression and pFSC048 containing the mFcγRI-CD3ζ construct were transformed into *E. coli* DH5α and cultured in LB media supplemented with the appropriate antibiotic. The plasmid DNA was extracted using the Qiagen QiaFilter Plasmid Midi and Maxi Kit (Qiagen, Germantown, MD) and linearized by restriction enzyme digestion to increase the efficiency of chromosomal integration into Jurkat cells. Plasmid pFSC005 was linearized by restriction enzyme digestion with SspI; plasmid pFSC048 was linearized by restriction enzyme digestion with PacI. Linearized plasmid DNA was purified using the Wizard® SV Gel and PCR Clean-up Kit (Promega, Madison, WI) in preparation for transfection into Jurkat cells.

#### Generation of Aequorin Expressing Platform Cells

Jurkat T cells were obtained from ATCC (Manassas, VA) and cultured in Complete RPMI 1640 medium following recommended ATCC guidelines. The cells were transfected with purified linear pFSC005 using the Nucleofector® 2b electroporator (Lonza, Morristown, NJ), following the Lonza Amaxa® Cell Line Nucleofector® Kit V optimized transfection protocol for Jurkat Clone E6-1 cells. Each transfection was performed with 4 μg of linearized DNA, using Lonza Program X-005 for maximum transfection efficiency. After transfection, cells were incubated at room temperature in a 12 well plate for 20 minutes before the addition of culture medium. The cell-containing plate was then incubated at 37°C, 5% CO_2_ for 24 hours, after which cells were centrifuged and resuspended in fresh culture medium. The Jurkat/hEF-1α-Aeq (Platform) cells were gradually expanded to 30 mL in Complete RPMI 1640 medium and cultured for 1 week until the cell viability exceeded 90%. Cells were then transferred to medium containing selection antibiotics (0.5 mg/mL G418) and cultured under selection for 2-3 weeks until cell viability recovered to at least 90%.

#### Generation of SPAR mFcγRI-CD3ζ Mixed Population Cells

Platform cells (Jurkat/hEF-1α-Aeq) were cultured under 0.5 mg/mL G418 selection as described above and transfected with linearized pFSC048 using the Lonza protocols described above. After transfection, cells were incubated in a 12 well plate at room temperature for 20 minutes before the addition of culture medium. The cell-containing plate was then incubated at 37°C, 5% CO_2_ for 24 hours, after which cells were centrifuged and resuspended in fresh culture medium. The Jurkat/hEF-1α-Aeq/mFcγRI-CD3ζ mixed population (SPAR mFcγRI-CD3ζ) cells were gradually expanded to 30 mL in Complete RPMI 1640 medium and cultured for 1 week until the cell viability exceeded 90% prior to the addition of selection antibiotics. Blasticidin was then added to a final concentration of 3 μg/mL to select for cells with chromosomal integration of mFcγRI-CD3ζ. Cells were cultured in Complete RPMI 1640 medium supplemented with 3 μg/mL blasticidin for 2-3 weeks to allow selection to occur and cell viability to recover to at least 90% before verification tests were performed.

#### Verification of mFcγRI-CD3ζ receptor expression and generation of SPAR Jurkat P5G7 cells by flow cytometry

The SPAR mFcγRI-CD3ζ cells were analyzed by flow cytometry to verify expression of the mFcγRI-CD3ζ receptor. Cells were washed with a FACS buffer containing FreeStyle™ 293 Expression Medium (Thermo Fisher Scientific) with 2% BSA and incubated with 15 µg/mL primary antibody, ChromPure Mouse IgG whole molecule (Jackson ImmunoResearch, West Grove, PA), for 30 minutes on ice. Samples were then washed twice with cold FACS buffer and incubated with 15 µg/mL secondary antibody, Alexa Fluor® 647 AffiniPure Goat Anti-Mouse IgG (Jackson ImmunoResearch), for 30 minutes on ice. After two additional washes in FACS buffer, the samples were run on a FACSAria II flow cytometer and analyzed using FlowJo software. A negative control sample of SPAR mFcγRI-CD3ζ cells stained with only secondary antibody was included to account for non-specific binding of the Alexa Fluor® 647 AffiniPure Goat Anti-Mouse IgG. An unstained sample of SPAR mFcγRI-CD3ζ cells was used for cell characterization and baseline gating. The clonal line SPAR Jurkat P5G7 was generated through single cell sorting from the top 10% of mFcγRI-CD3ζ expressing SPAR mFcγRI-CD3ζ mixed population cells. SPAR Jurkat P5G7 cells were expanded and analyzed for mFcγRI-CD3ζ receptor expression using the method described above.

#### Verification of Aequorin and mFcγRI-CD3ζ receptor transfection and expression via antibiotic selection and PCR

Jurkat/hEF-1α-Aeq platform cells were cultured under antibiotic selection in RPMI 1640 supplemented with 25 mM dextrose, 10 mM HEPES, 1 mM sodium pyruvate, 10% FBS, 1% Pen-Strep (hereinafter referred to as Complete RPMI 1640 medium) containing 0.5 mg/mL G418 sulfate to select for chromosomal integration of pEF1-Aeq. SPAR Jurkat P5G7 cells were cultured under antibiotic selection in growth medium containing 3 µg/mL blasticidin to select for chromosomal integration of the mFcγRI-CD3ζ construct. Cells were cultured under selection for 2-3 weeks until the cell viability recovered to at least 90% before further analysis. Genomic DNA was extracted from SPAR Jurkat P5G7 cells using the Qiagen DNeasy® Blood & Tissue Kit. Chromosomal gene insertion was verified by PCR to confirm the fusion of mFcγRI with CD3ζ. Primer FSC-1 targeted a sequence within FcγRI and primer FSC-2 targeted a sequence between CD3ζ and the IRES sequence. The PCR product was analyzed by gel electrophoresis to confirm the presence of the correct band size in the SPAR Jurkat P5G7 sample and the absence of a band in the negative control Platform cell genomic DNA sample.

#### Verification of Aequorin expression by generation of light signal

Platform or SPAR Jurkat P5G7 cells were resuspended to a concentration of 1.1 x 10^6^ cells/mL in RPMI 1640 supplemented with 25 mM dextrose, 10 mM HEPES, 1 mM sodium pyruvate, 10% Ultra Low IgG FBS, 1% Pen-Strep, and 0.1% Pluronic F68 (hereinafter referred to as Charging medium) in 30 mL conical tubes. Cells were charged by addition of the luminescent substrate coelenterazine-h to a final concentration of 1.5 µM. The cell suspension was protected from light and incubated at room temperature with shaking (60 RPM) for 18-24 hours to allow the coelenterazine-h to enter the cells and bind to expressed aequorin. Light signal generation was evaluated by adding 180 µL of charged Jurkat platform cells (400,000 cells/180 µL) to 30 µL of 0.61 mM digitonin in a 1.5 mL microcentrifuge tube that had been pre-inserted in the luminometer reader (GloMax, Promega). The signal was recorded for 1 minute. The assay was repeated using the same concentration of charged Jurkat parental cells, uncharged Jurkat parental cells, and uncharged Jurkat platform cells as negative controls.

#### Generation of serotype-specific mouse monoclonal antibody against *E. coli* O111 LPS

Serotype-specific mouse monoclonal antibody against *E. coli* O111 LPS was generated by Precision Antibody (Columbia, MD) using antigen prepared and supplied by this laboratory. Briefly, initial immunizations were performed by injecting mice with a deactivated bacteria cell wall preparation until a robust immune response was achieved. LPS was then extracted from isolated bacteria cell wall preparation in a process that enriched LPS and eliminated other soluble components of bacteria such as nucleic acids and cytoplasmic proteins (24). Following the initial immunization, the purified LPS was then used to boost mice 2-3 times before their spleen cells were fused with myeloma. At 10-12 weeks after the initial injection, final hybridoma clones were collected and screened by ELISA using LPS from *E. coli* O111 and other bacterial pathogens.

#### ELISA detection of antibodies

A 96-well microtiter plate was coated with *E. coli* O111 LPS at 4°C overnight, then washed to remove unbound antigen followed by blocking with 5% w/v BSA (or nonfat dry milk) at room temperature for 1 hour. Serial dilutions of antibody samples, including standards, positive and negative controls, and unknowns were added to the plate. After incubating for 1hour the plate was washed three times and then incubated with HRP-conjugated detection antibody for 1 hour. The plate was washed and a 3,3’,5,5’-Tetramethylbenzidine substrate solution was added for color development. The reaction was stopped by 1M HCl and the plate was immediately read using a plate reader at absorbance wavelength of 450 nm.

### Detection of Bacterial Pathogens Using SPAR Jurkat P5G7 Cells

#### *E. coli* O111 Bacteria

Charged SPAR cells were centrifuged and resuspended to a concentration of 2.2 x 10^6^ cells/mL in charging medium. Prior to starting the assay, 180 µL of resuspended SPAR cells were incubated with mouse anti-*E. coli* O111 LPS IgG in a 1.7 mL microcentrifuge tube to a final antibody concentration of 9.8 µg/mL at room temperature for 10 minutes. To begin the assay, the antibody-coated SPAR cells were added to an overnight culture of *E. coli* O111 that was serially diluted to a concentration of 10,000 CFUs/mL (confirmed by plating and counting) and was already placed in the luminometer. The light signal was recorded immediately and continuously for 4 minutes. Additional controls included 1) testing without antibody, wherein 180 µL of the resuspended biosensor cells were added directly to the serially diluted *E. coli* O111 in a 1.7 mL microcentrifuge tube placed in the luminometer, and 2) testing without bacteria, in which 180 µL of the biosensors were added directly to 1.6 µL of 1.25 mg/mL mouse anti-*E. coli* O111 LPS IgG in a 1.7 mL microcentrifuge tube in the luminometer. All tests were performed in triplicate.

#### *E. coli* O111 LPS

Charged SPAR cells were centrifuged and resuspended to a concentration of 2.2 x 10^6^ cells/mL in charging medium. To begin the assay, 180 µL of resuspended charged SPAR cells were mixed with mouse anti-*E. coli* O111 LPS IgG in a 1.7 mL microcentrifuge tube to a final antibody concentration of 9.8 µg/mL and then added to 30 µL of *E. coli* O111 LPS in a second 1.7 mL microcentrifuge tube that was already placed in the luminometer. The light signal was recorded immediately and continuously for 4 minutes. The assay was repeated with *E. coli* O157 LPS as negative control. Additional controls included 1) testing without antibody, wherein 180 µL of the resuspended biosensor cells were added directly to *E. coli* O111 LPS in a 1.7 mL microcentrifuge tube placed in the luminometer, and 2) testing without LPS, in which 180 µL of the biosensors were added directly to 1.6 µL of 1.25 mg/mL mouse anti-*E. coli* O111 LPS IgG in a 1.7 mL microcentrifuge tube in the luminometer. All tests were performed in triplicate.

#### *P. aeruginosa* Bacteria

Charged SPAR cells were centrifuged and resuspended to a concentration of 1.6 x 10^7^ cells/mL in charging medium. Prior to starting the assay, 90 µL of resuspended charged SPAR cells were mixed with mouse anti-*P. aeruginosa* IgG to a final antibody concentration of 1.5 µg/mL. To begin the assay, the antibody-coated cells were added to 30 µL of an overnight culture of *P. aeruginosa* that was serially diluted to a concentration of 1000 CFUs/mL (confirmed by plating and counting) in a 1.7 mL microcentrifuge tube already placed in the luminometer. The light signal was recorded immediately and continuously for 8 minutes. As a negative control test, the assay was repeated using an overnight culture of *S. enteritidis* serially diluted to a similar concentration as the *P. aeruginosa*. Additional controls included 1) testing without antibody, wherein 90 µL of resuspended charged SPAR cells was added directly to a 30 µL aliquot of diluted *P. aeruginosa*, and 2) testing in the absence of bacteria, in which 90 µL of SPAR cells were added to 1.8 µL of 0.1 mg/mL mouse anti-*P. aeruginosa* IgG in a microcentrifuge tube already placed in the luminometer. All tests were performed in triplicate.

#### *E. coli* O157 Bacteria

Charged SPAR cells were centrifuged and resuspended to a concentration of 4.4 x 10^6^ cells/mL in charging medium. A solution of coupled antibodies was prepared by mixing mouse anti-Human IgG (mouse IgG cross-reacted with goat IgG) with goat anti-*E. coli* O157 LPS IgG to concentrations of 0.4 mg/mL and 0.1 mg/mL, respectively, in RPMI 1640 with 1% BSA. The antibody solution was incubated for 30 minutes before use. Prior to starting the assay,180 µL of resuspended SPAR cells were incubated with 6 µL of the coupled antibody solution in a 1.7 mL microcentrifuge tube at room temperature for 10 minutes. The final antibody concentrations were 11.4 µg/mL and 2.9 µg/mL respectively. To begin the assay, the antibody-coated SPAR cells were added to 30 µL of an overnight culture of *E. coli* O157 that was serially diluted to 10,000 CFUs/mL (confirmed by plating and counting). The light signal was recorded immediately and continuously for 8 minutes. As a negative control, the assay was repeated using a similarly diluted overnight culture of *E. coli* O111.As an additional control, testing was performed without antibody by adding 180 µL of resuspended charged SPAR cells directly to a 30 µL aliquot of the diluted *E. coli* O157 culture. All tests were performed in triplicate.

## Results

### Design of Gene Constructs

To generate the SPAR cell modified TCR complex (Figure 1, **lower panel**), the CD3ζ subunit of the TCR complex was genetically engineered to be expressed as a fusion protein where the extracellular domain of CD3ζ was fused with FcγRI. A short GS linker was genetically introduced to separate the antibody binding domain FcγRI from the signal-transducing protein element CD3ζ, thus ensuring proper folding of the engineered protein fragments. The CD3ζ signal peptide sequence was used to specify cell surface expression of the FcγRI-linker-CD3ζ fusion protein.

**Figure 1.**
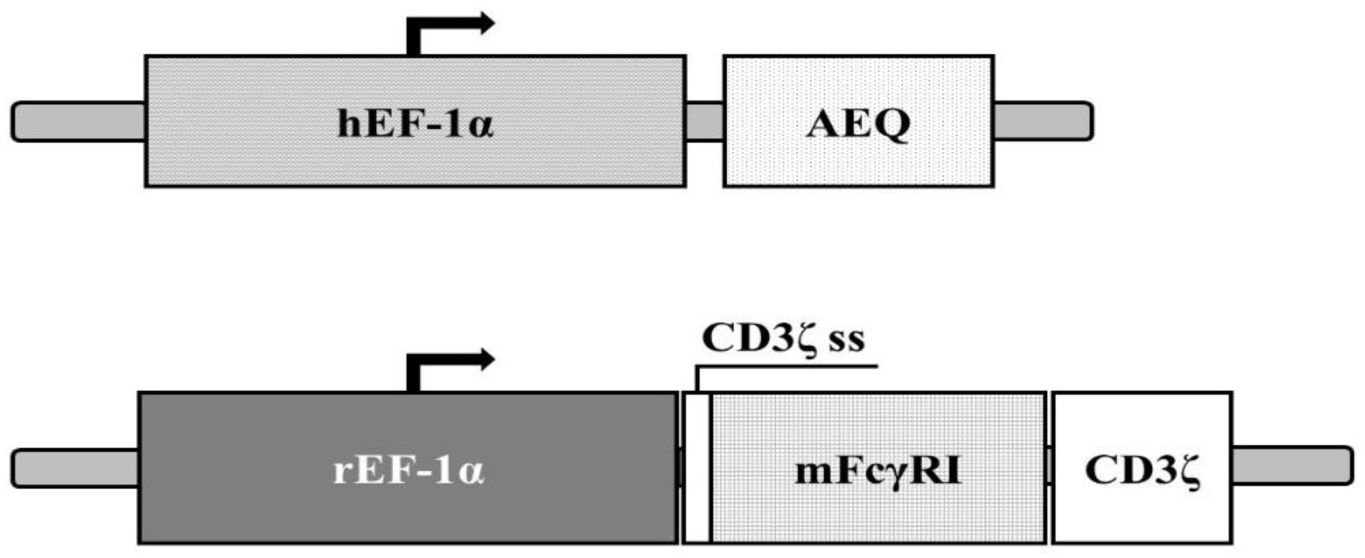
Graphic depiction of the SPAR Jurkat P5G7 gene constructs. Upper panel: The luminescent reporter enzyme Aequorin (AEQ) gene fused with the human EF-1*α* constitutive promoter. Lower panel: The receptor construct mFcγRI-CD3ζ linked to the CD3ζ signal sequence and rat EF-1*α* constitutive promoter.

The cytoplasmic aequorin luminescent reporter system (Figure 1, **upper panel)** was designed for high level expression by fusion of the human elongation factor 1α-subunit promoter with a codon-optimized, commercially synthesized aequorin gene (DNA2.0).

When expressed, the modified TCR complex (Figure 2 B) was designed to conserve the majority of the native TCR complex conformation, retaining all ten of the immunoreceptor tyrosine-based activating motifs (ITAMS) expressed by the unmodified TCR complex (Figure 2 A), thereby preserving maximum signaling ability. The mFcγRI IgG binding site is accessible on the cell surface and can bind to the Fc region of murine IgGs.

**Figure 2.**
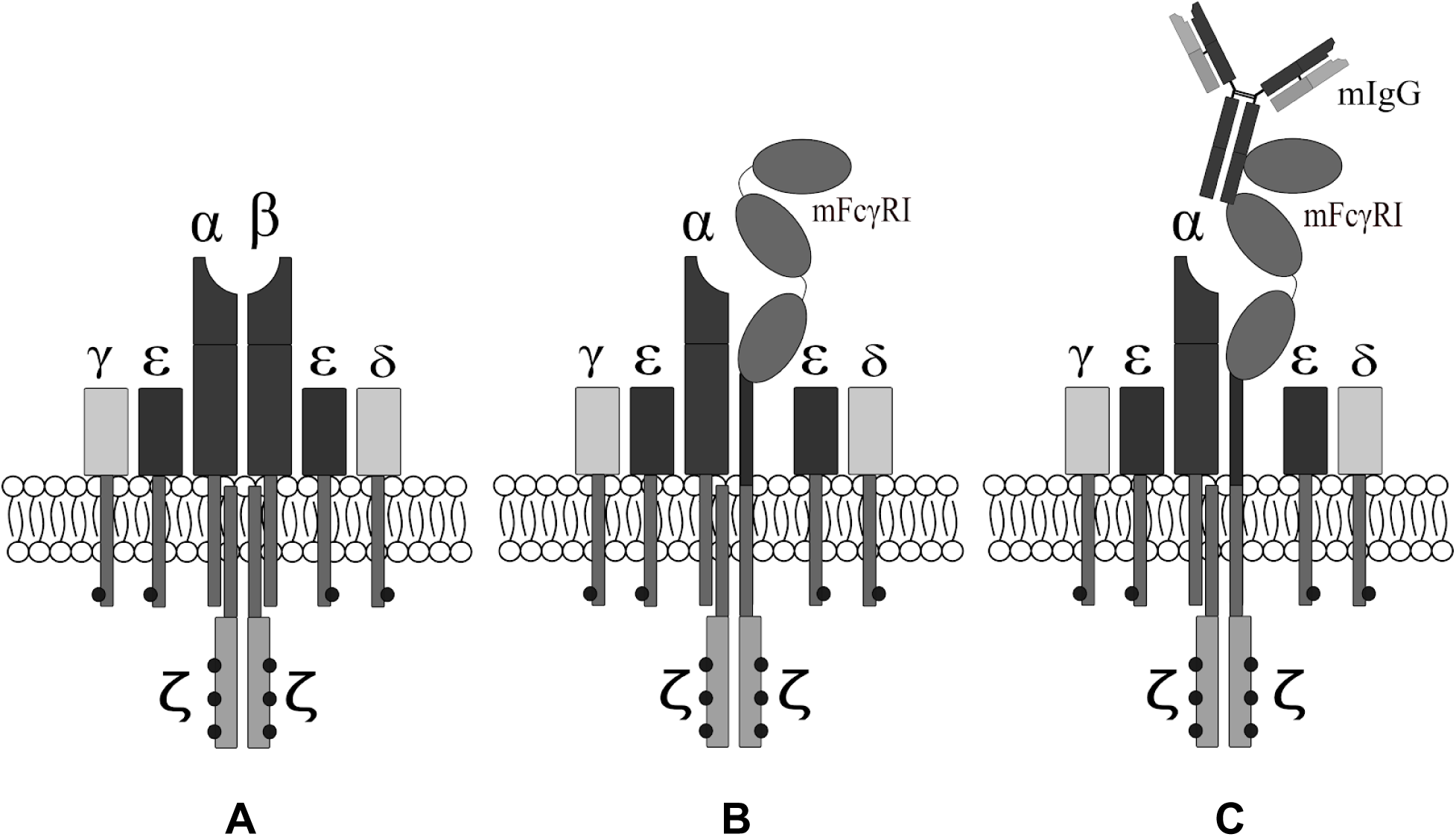
Schematic depiction of the SPAR cell receptor design. The system was generated through modification of the TCR complex. (**A**) Unmodified T-cell receptor complex; (**B**) Mouse FcγRI fused to CD3*ζ* (mFcγRI-CD3*ζ*) and (**C**) Mouse IgG bound to mFcγRI-CD3*ζ*.

### Verification of FcγRI-CD3ζ receptor expression

Expression of the mFcγRI-CD3ζ receptor in SPAR Jurkat P5G7 cells was confirmed by flow cytometry using a two-step staining process. SPAR Jurkat P5G7 cells were incubated with the primary antibody ChromPure mouse IgG whole molecule, which binds to the mFcγRI-CD3ζ receptor, followed by staining with the secondary antibody Alexa Fluor® 647 AffiniPure Goat Anti-Mouse IgG. The level of receptor expression was measured by the amount of fluorescence detected by a BD FACSAria II flow cytometer equipped with a 633 nm laser and a 670/30 bandpass filter. The baseline fluorescence level of SPAR Jurkat P5G7 cells determined by using an unstained cell sample. Negative control gating was established using SPAR Jurkat P5G7 cells stained with Alexa Fluor® 647 AffiniPure Goat Anti-Mouse IgG only. The negative control sample showed minimal non-specific binding of the secondary antibody when compared to the unstained sample (Figure 3). SPAR Jurkat P5G7 cells stained with ChromePure Mouse IgG Whole Molecule and Alexa Fluor® 647 AffiniPure Goat Anti-Mouse IgG produced a significant fluorescent shift compared to the negative control, indicating the presence of the mFcγRI-CD3ζ receptor on the SPAR Jurkat P5G7 cell surface as well as the ability of the receptor to bind mouse IgG antibodies (Figure 3).

**Figure 3.**
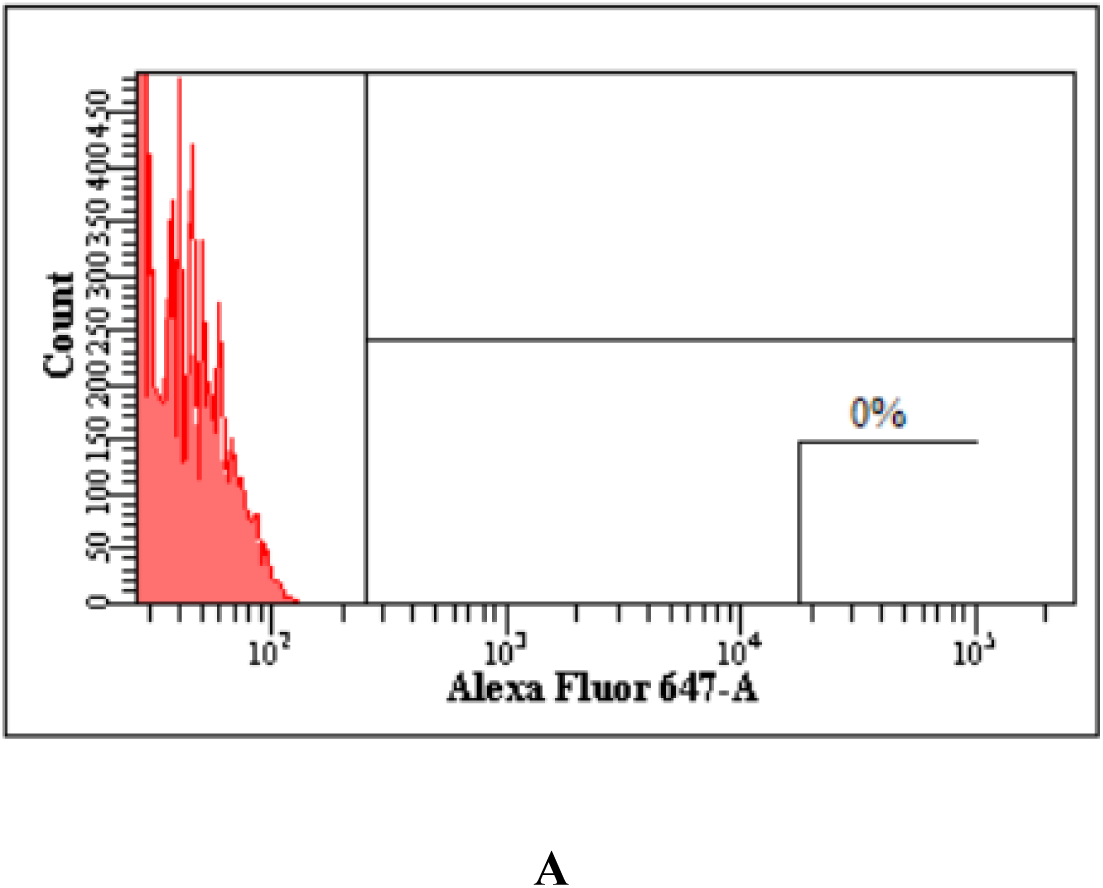
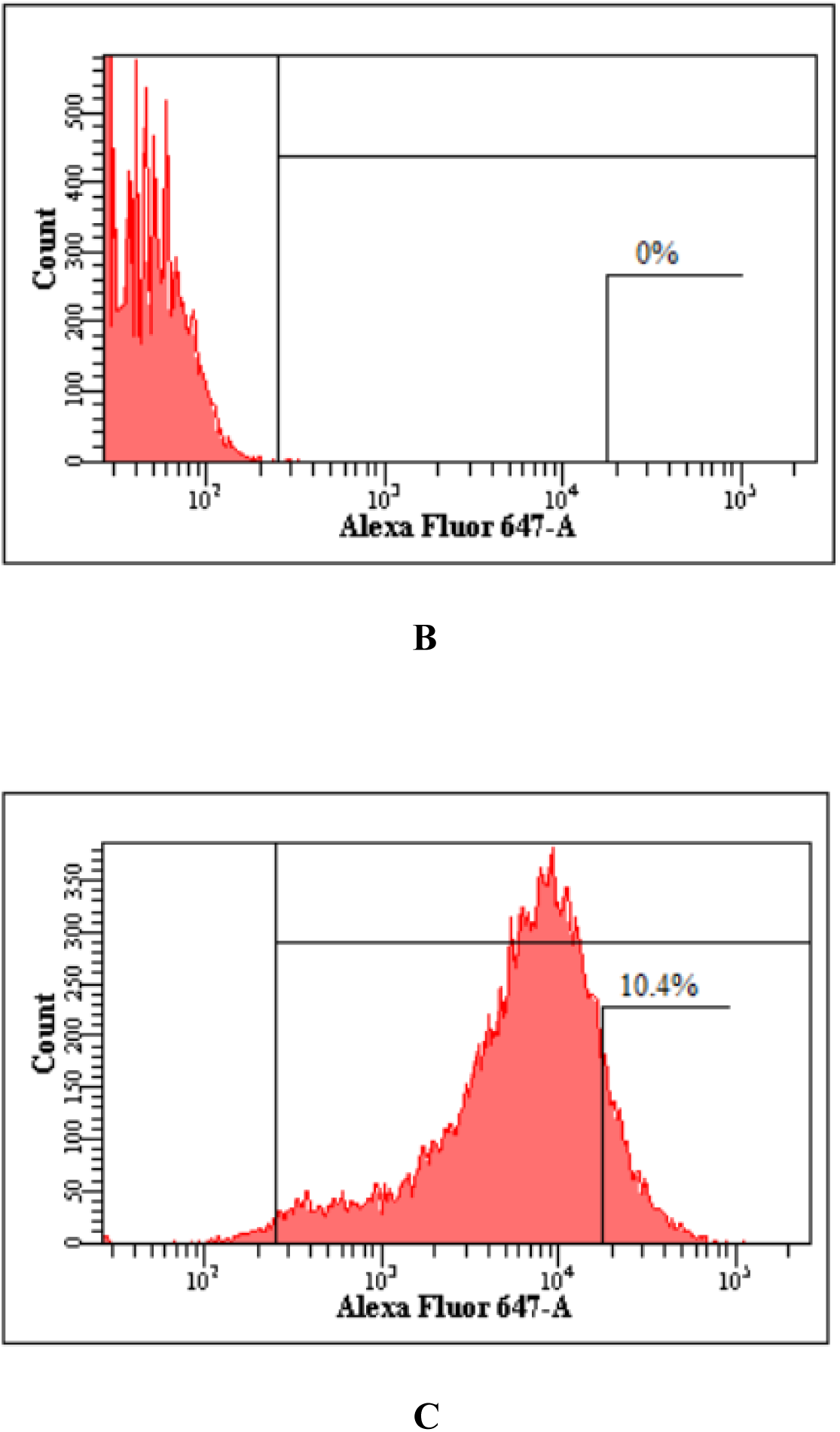
Flow cytometric verification of mFcγRI-CD3ζ receptor expression in SPAR Jurkat P5G7 cells stained with ChromePure Mouse IgG Whole Molecule and Alexa Fluor® 647 AffiniPure Goat Anti-Mouse IgG. **A**) Unstained SPAR Jurkat P5G7 cells; **B**) SPAR Jurkat P5G7 cells stained with Alexa Fluor® 647 AffiniPure Goat Anti-Mouse IgG (secondary Ab) alone to account for non-specific binding of the secondary antibody; **C**) SPAR Jurkat P5G7 cells stained with ChromePure mouse IgG whole molecule (primary Ab) and Alexa Fluor® 647 AffiniPure Goat Anti-Mouse IgG (secondary Ab).

### Verification of mFcγRI-CD3ζ receptor transfection by PCR

Successful, stable transfection was verified via PCR and gel electrophoresis (Figure 4). Chromosomal gene insertion of the mFcγRI-CD3ζ fusion construct in SPAR Jurkat P5G7 cells was verified by PCR (Figure 4). The PCR product was analyzed by gel electrophoresis. Results confirmed the presence of the correct band size (0.690 kb) in the SPAR Jurkat P5G7 sample and the absence of a band in the negative control Jurkat platform cell sample.

**Figure 4.**
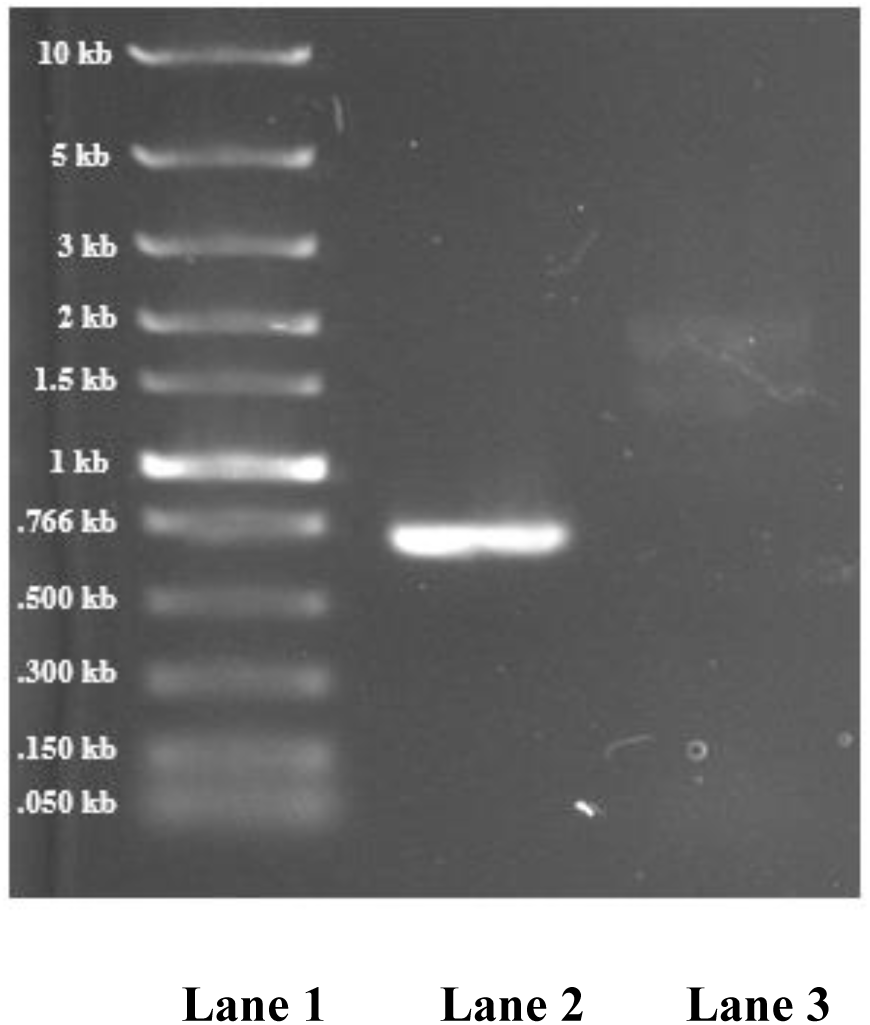
Verification of chromosomal integration of the mFcγRI-CD3ζ fusion construct in SPAR Jurkat P5G7 cells by PCR and gel electrophoresis. **Lane 1**: Standard ladder. **Lane 2**: SPAR Jurkat P5G7 Genomic DNA (band size 0.690 kb). **Lane 3**: Platform cell genomic DNA (negative control).

### Demonstration of aequorin reporter system function

To verify that the expressed aequorin reporter system is functional in the Jurkat platform cells, we evaluated the ability of charged platform cells to generate a light signal in response to digitonin treatment, which provides a receptor-independent stimulation of aequorin. Only charged platform cells produced a dramatic luminescent signal following the addition of digitonin (Figure 5). No significant signal generation was noted after treatment of uncharged Jurkat platform cells, uncharged Jurkat parental cells or charged Jurkat parental cells, indicating that the signal recorded was specific and dependent upon both the expression of aequorin and the prior charging of the reporter system by the addition of coelenterazine-h.

**Figure 5.**
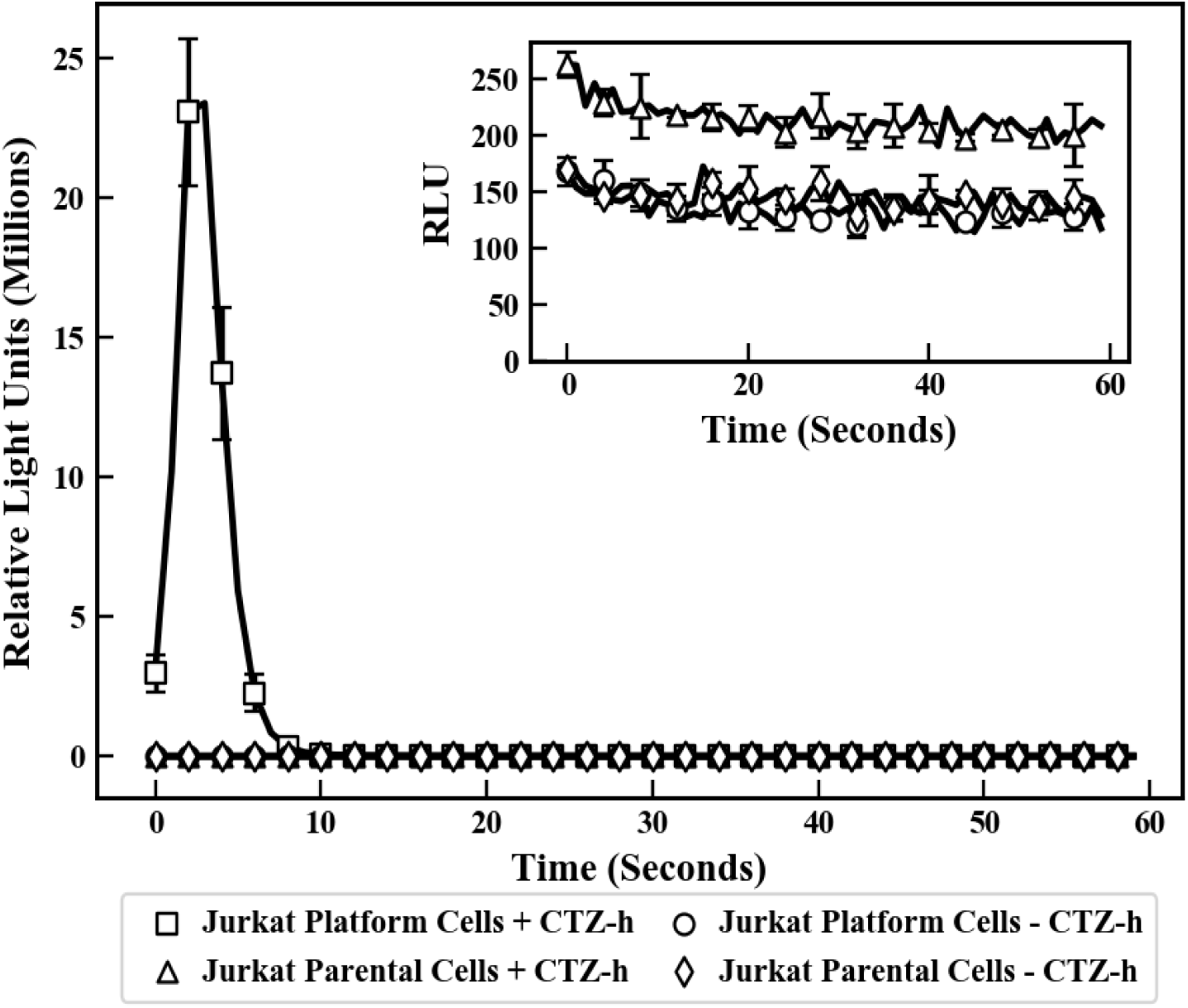
Receptor-independent verification of aequorin expression by generation of a light signal. 180 µL of resuspended Jurkat platform cells charged by incubation with coelenterazine-h (+CTZ-h) were mixed with 30 µL of 0.61 mM digitonin to generate a light signal. Charged Jurkat parental cells, uncharged (-CTZ-h) Jurkat platform cells, and uncharged Jurkat parental cells were tested as negative controls. All cells were tested at 2.2 x 10^6^ cells/mL. Each data point is a mean of triplicates and error bars represent one standard deviation from the mean. Inset, chart showing detail of the response demonstrated by charged Jurkat parental cells, uncharged Jurkat platform cells, and uncharged Jurkat parental cells.

### Validation of SPAR Jurkat P5G7 cell signal transduction

The results reported above demonstrate that SPAR Jurkat P5G7 cells express both the FcγRI-CD3ζ cell surface receptor and an intact aequorin reporter system. The clinical utility of SPAR Jurkat P5G7 cells for pathogen detection depends upon the functional linkage of these two properties. Thus, SPAR Jurkat P5G7 cells were exposed to an antibody specific for the human CD3ε subunit of the TCR complex and tested to evaluate the ability of the resulting receptor aggregation to trigger signal transduction and subsequently activate the reporter system, demonstrated by the emission of detectable light. The binding of human CD3ε-specific antibody (Biolegend, San Diego, CA) to charged SPAR Jurkat P5G7 cells resulted in the generation of a strong light signal measurable within 50 seconds and peaking at 120 seconds (data not shown). No signal was detected when an antibody specific for the B lymphocyte antigen CD19 (25) was used as negative control. These results verify the cell surface expression of the TCR complex, the presence of an intact and functional signal transduction pathway, and the ability of the aequorin detection system to specifically respond to Ca^2+^ release.

### Generation of serotype-specific monoclonal antibody against *E. coli* O111 LPS

The lipopolysaccharide (LPS) exposed on the cell surface and localized to the outer layer of the membrane of Gram-negative bacteria has been shown to be an important virulence factor (26). Intact bacterial lipopolysaccharides (LPS) are macromolecules made up of three structural components: Lipid A, polysaccharide chain, and O-antigen. Distinctive O-antigen structures have been used to classify serogroups to *E. coli*, *S. enteritidis*, and *V. cholerae* (27); thus, *E. coli* O111 LPS was used to generate this serotype-specific monoclonal antibody. The use of a cell wall preparation dramatically reduced the time and labor required for hybridoma screening and shortened the timeline for monoclonal antibody production. ELISA evaluation of antibody binding to a variety of pathogens (Figure 6) demonstrated high specificity to *E. coli* O111 and indicated no cross-reactivity with alternate strains of *E. coli* (O157 or O26) or to different pathogens.

**Figure 6.**
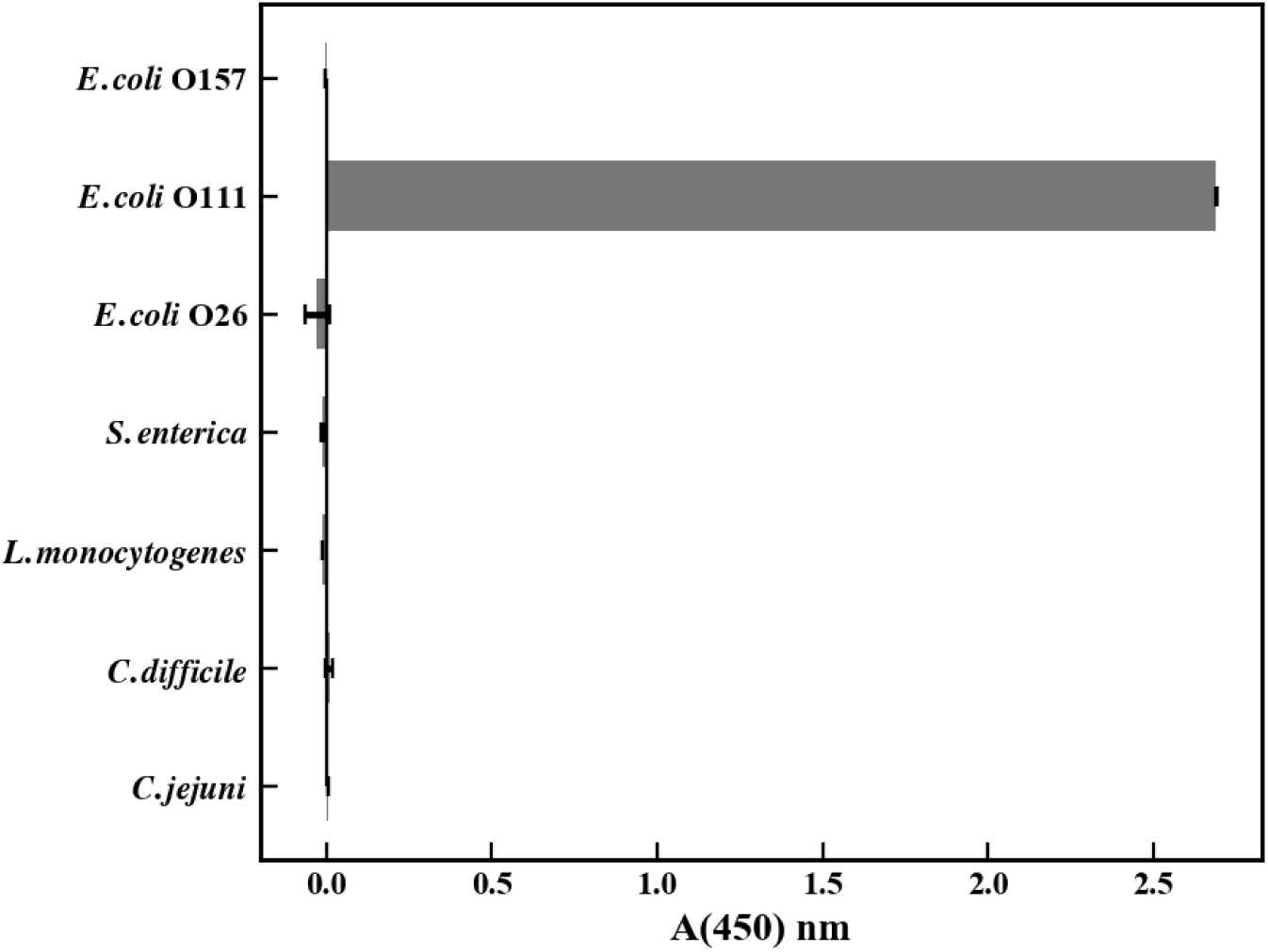
Specificity of the monoclonal antibody generated against *E. coli* O111 LPS as demonstrated by ELISA response when analyzed against a variety of pathogens.

### SPAR Jurkat P5G7 cell detection of *E. coli* O111 LPS

The ELISA results described above document the specificity of the monoclonal antibody generated against *E. coli* O111; thus, we evaluated the ability of SPAR cells complexed with this antibody to detect purified *E. coli* O111 LPS. Figure 7 depicts the signal generated by charged, antibody-complexed SPAR cells in response to the addition of purified *E. coli* O111 LPS. The initiation of a significant light signal was observed within 30 seconds of antigen addition and peaked at approximately 60 seconds. The addition of purified *E. coli* O157 LPS was used as control and produced no signal response, demonstrating the specificity of detection.

**Figure 7.**
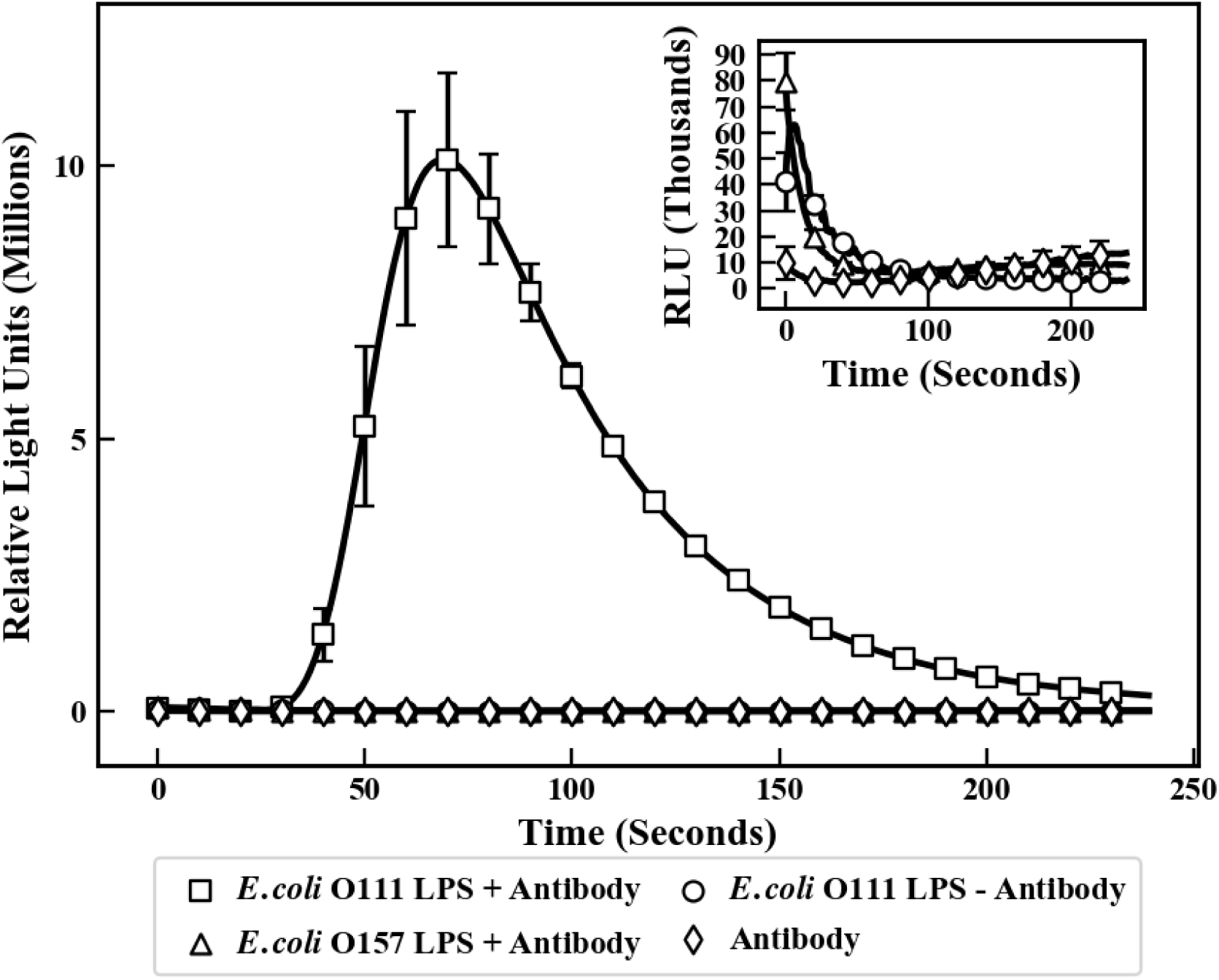
Detection of *E. coli* O111 LPS using 400,000 SPAR Jurkat P5G7 cells in 180 µL medium. Charged SPAR cells were mixed with 9.8 μg/mL mAb against *E. coli* O111 LPS before adding to 30 µL of 142 µg/mL *E. coli* O111 LPS to generate a light signal. None of the three controls (SPAR cells + Antibody + *E. coli* O157 LPS, SPAR cells + Antibody, and SPAR cells + *E. coli* O111 LPS) generated a light signal. Inset: a chart showing greater detail of the minimal response demonstrated by the control samples. Each data point is a mean of triplicates and error bars represent one standard deviation from the mean.

### SPAR Jurkat P5G7 cell detection of bacterial samples

To demonstrate the clinical utility of SPAR cells in pathogen detection, charged SPAR cells were customized by the addition of a variety of specific antibodies and incubated with corresponding bacterial samples derived from live culture. Detection ability and specificity were evaluated as above, using signal generation as read by the GloMax luminometer. Results are depicted below in Figure 8 (*E. coli* O111), Figure 9 (*P. aeruginosa*), and Figure 10 (*E. coli* O157).

**Figure 8.**
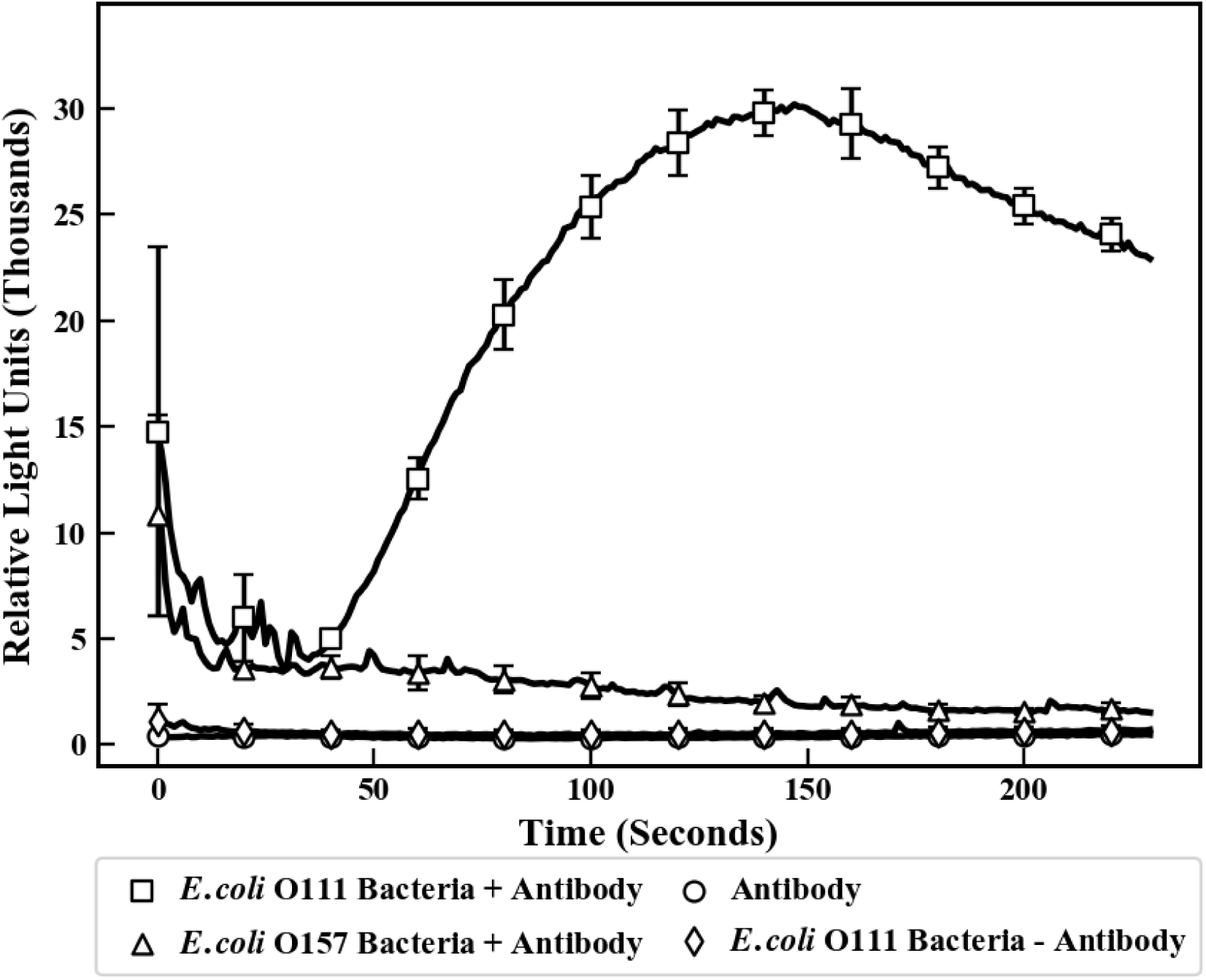
Detection of *E. coli* O111 bacteria using SPAR Jurkat P5G7 cells (2.2 x 10^6^ cells/mL). 180 µL of resuspended SPAR cells were coated with 9.8 μg/mL monoclonal antibody against *E. coli* O111 LPS before mixing with 10,000 CFUs/mL of serially diluted *E. coli* O111 bacterial culture. None of the three controls (SPAR cells + antibody + *E. coli* O157 bacteria, SPAR cells + antibody, and SPAR cells + *E. coli* O111 bacteria) generated detectable light signal. Each data point is a mean of triplicates and error bars represent one standard deviation from the mean.

**Figure 9.**
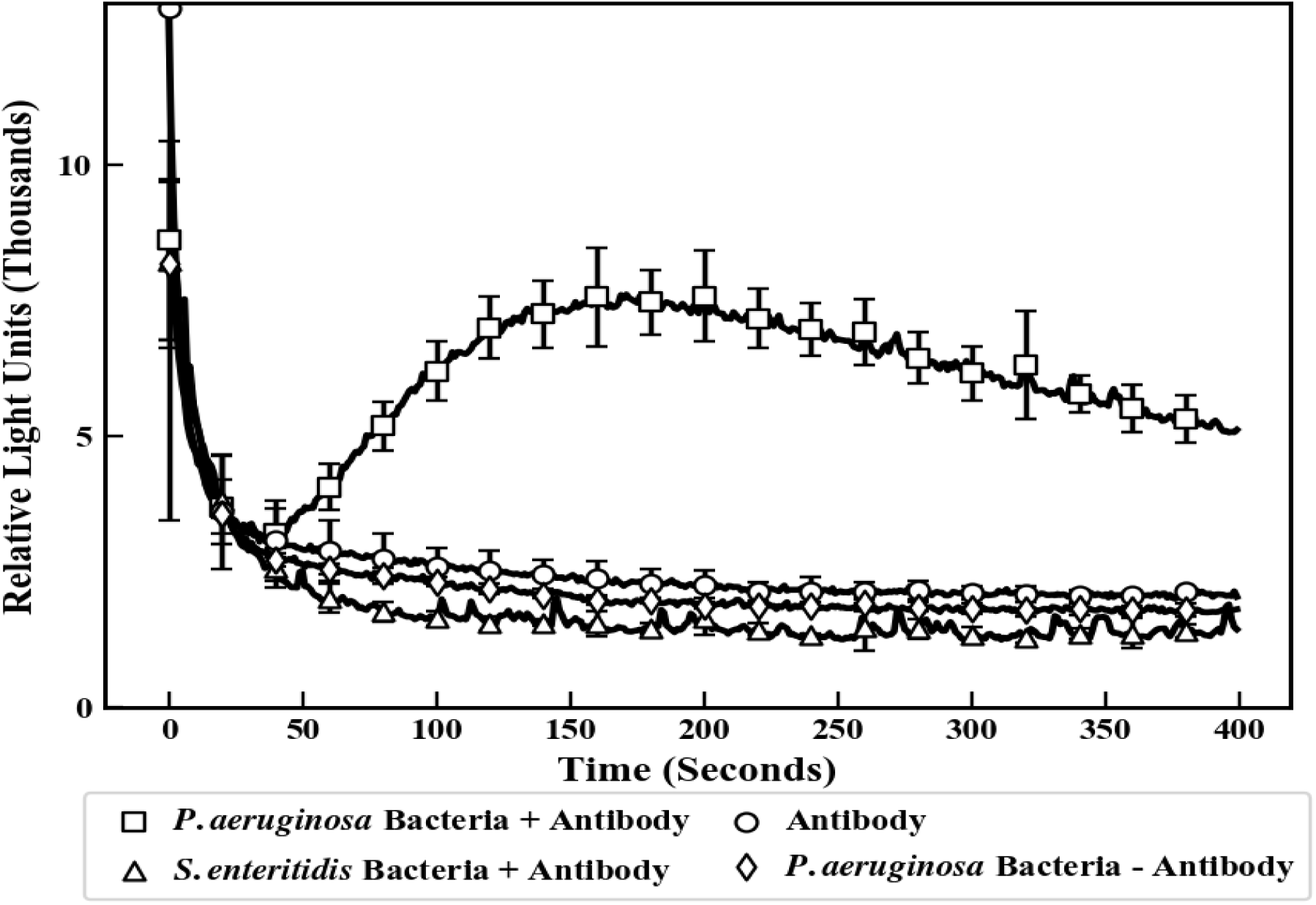
Detection of *P. aeruginosa* using SPAR Jurkat P5G7 cells (2.2 x 10^6^ cells/mL). 180 µL of resuspended SPAR cells were coated with 9.8 μg/mL. Charged cells were pre-loaded with a monoclonal antibody against *P. aeruginosa* before mixing with serially diluted culture samples containing the target pathogen. None of the three controls (non-specific *S. enteritidis* + antibody, *P. aeruginosa* without antibody, and antibody alone without target) generated detectable light signal. Each data point is a mean of triplicates and error bars represent one standard deviation from the mean. The detector sensitivity for *P. aeruginosa* was 1000 CFUs/mL.

**Figure 10.**
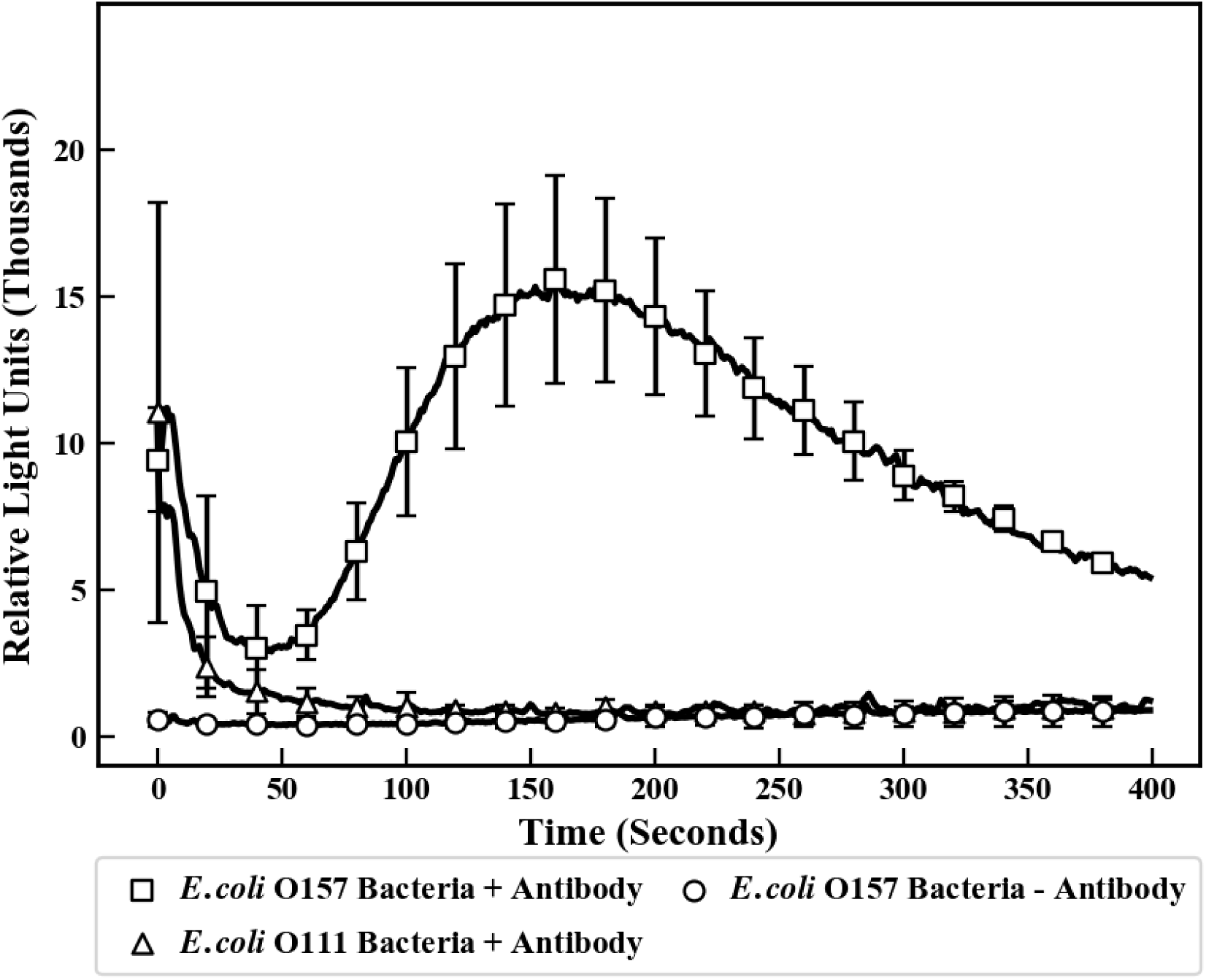
Detection of *E. coli* O157 bacteria using the SPAR Jurkat P5G7 cells (2.2 x 10^6^ cells/mL) in a coupled antibody system. 90 µL of the resuspended SPAR Jurkat P5G7 cells were coated with mouse anti-human IgG coupled with goat anti-*E. coli* O157 before mixing with serially diluted *E. coli* O157 bacteria. No signal was generated by negative controls (*E. coli* O111 bacteria + coupled antibodies, and *E. coli* O157 bacteria without antibodies). Error bars represent one standard deviation from the mean of three runs. The detector sensitivity was 10,000 CFUs/mL.

## Discussion

We have engineered two plasmids that, when concurrently stably transfected into Jurkat cells, generate a living biosensor cell platform capable of the rapid and specific detection of a variety of molecules of interest. In this study, we demonstrate that the SPAR platform can be used to reliably detect bacterial pathogens in samples generated from overnight bacterial culture. The detection assay is complete and can be read within minutes, utilizes standard laboratory equipment, and requires no advanced training of personnel.

The SPAR platform biosensor cells are designed to stably express both the modified mFcγRI-CD3ζ universal cell surface receptor and an aequorin reporter system. Fc receptors are expressed on the surface of hematopoietic cells and function as essential components of the antibody-mediated immune response and thus have been well characterized (28, 29). Each type of Fc receptor reacts specifically with a corresponding class of immunoglobulin: FcγRI receptors have been demonstrated to bind to their cognate antibodies with high affinity (28, 30, 31) though the affinity of the mFcγRI receptor for mouse antibodies varies with antibody subclass (32, 33). Mouse IgG2a antibodies have the highest affinity, with dissociation constants (K_D_) in the nanomolar range (34). Native FcγRs consist of a ligand-binding FcγR α-chain expressed on the cell surface, and a γ chain dimer which functions in signal transduction and bears immunoreceptor tyrosine-based activating motifs (ITAMS) (35).

PCR results confirmed expression of the modified mFcγRI-CD3ζ receptor in SPAR Jurkat P5G7 cells; additionally, we verified that the receptor is expressed on the cell surface, a property required for receptor function (32). Only SPAR cells shown to express the engineered receptor are capable of binding mouse IgG, which was subsequently complexed with Alexa Fluor® 647 AffiniPure Goat Anti-Mouse IgG and visualized via flow cytometry.

Prior research has shown that the signal transduction pathways of Jurkat cells are intact and functional: antigen binding to TCR-positive Jurkat cells triggers rapid intracellular Ca^2+^ release (35, 36). This intracellular Ca^2+^ release is detectible using the well-characterized aequorin reporter system (19, 21, 37). SPAR cells were therefore engineered to express aequorin, activated via the addition of coelenterazine-h, as a mechanism to visibly detect antigen binding and subsequent signal transduction. Aequorin expression by SPAR cells was verified by flash testing: the free aequorin discharged by digitonin-lysed SPAR cells generates a light signal and does so only when SPAR cells are charged by prior incubation with coelenterazine.

Our results further demonstrate that engineered SPAR cells effectively detect bacterial pathogens: this detection can be customized by the addition of pathogen-specific antibodies. Binding of a monoclonal antibody generated against *E. coli* O111 LPS to charged SPAR cells detects both isolated *E. coli* O111 LPS and intact *E. coli* O111 sampled from overnight bacterial culture. Specificity of detection is demonstrated by an absence of signal when the charging step is omitted, and by a lack of cross-reactivity to *E. coli* O157. Similarly, binding of a commercially available monoclonal antibody specific for *P. aeruginosa* detected the presence of *P. aeruginosa* in samples from overnight culture and indicated no cross-reactivity with *S. enteritidis*.

In cases where no commercially produced mouse antibody is available against the pathogen of interest, a sandwich technique is effective and has been used here for the detection of *E. coli* O157. SPAR platform cells were incubated with a mouse anti-human IgG demonstrated to be cross-reactive with goat IgGs and subsequently coupled to goat anti-*E. coli* O157. Addition to *E. coli* O157 sampled from overnight cultures specifically generated light signal; no cross-reactivity was evident upon addition to *E. coli* O111.

The generation of light signal reaches peak levels within 200 seconds regardless of the antibody employed. The brief light signal detected within the initial few seconds of bacterial assay is likely the result of damage incurred to the cell membrane of a small subset of SPAR cells during handling, thus allowing a receptor-independent stimulation of free aequorin similar to that depicted in Figure 5. This free aequorin is rapidly exhausted and does not contribute to the specific light signal generated in response to antigen recognition.

Repeat experiments have demonstrated that reaction efficiency appears to be independent of the order of reactant addition. SPAR cell number was optimized during the course of this study. It was found that a concentration of 1.8 x 10^7^ cells/mL, as compared to the initial concentration of 2.2 x 10^6^ cells/mL, improves detection sensitivity. This system is extremely sensitive and able to detect small amounts of target analyte: the biological amplification provided by the SPAR cell circumvents the need for time-consuming sample enrichment steps characteristic of alternative detection methods.

In conclusion, the results presented in this paper demonstrate that the modified TCR complex engineered in this laboratory is expressed on the SPAR cell surface, is functional, and when complexed with the appropriate antibody responds appropriately to its cognate antigen. The signal transduction pathway is triggered and intracellular Ca^2+^ levels increase due to the rapid release of Ca^2+^ from the endoplasmic reticulum and the subsequent opening of calcium release-activated channels allowing Ca^2+^ influx from the medium. This rapid increase in cytosolic Ca^2+^ activates the aequorin system constitutively expressed in the SPAR cell cytoplasm, generating a robust light signal that can be detected within minutes using standard laboratory instrumentation. This system uses a single, stable cell line—the SPAR cell—which is programmable through the addition of specific antibodies, whether commercially available or generated in-house. The SPAR platform thus provides a rapid, sensitive, and convenient method for the detection of a variety of pathogens.

## Acknowledgments

Michelle Pate, Research Assistant, Academic Research Center, Department of Chemical and Biomolecular Engineering, Ohio University, Athens Ohio, for assistance in flow cytometry.

The OU Innovation Center for technical support in growing hybridomas.

## Support

This research was supported by FSC/CytoSPAR and received no specific grant from any funding agency in the public or not-for-profit sectors.

## Affiliations/consulting agreements, etc.

Charles McBrairty and M. Russell Willams are founders of FSC/CytoSPAR and currently hold equity interests in FSC/CytoSPAR. Joseph D. Kittle is currently Chief Scientific Officer of FSC/CytoSPAR. Thomas Zupancic is Chief Scientific Officer of InfinixBio; Rich Brody is Vice President of Research. Joel S. Lwande, Shengwen Liang, Yuanyuan Tang, Jiangzhou Hua are employees of InfinixBio. Kyle McQuaid, Melissa Frenchmeyer, Yuanyuan Tang, Allison Neese, and Jiangzhou Hua are employees of DevLab.

## Statement of medical writer contribution

The authors thank Joan Breslin, PhD of CSSi LifeSciences, Glen Burnie, MD for providing medical writing support/editorial support which was funded by FSC/CytoSPAR, Easton PA in accordance with Good Publication Practice (GPP3) guidelines.

